# Edge-focused network-based approach: an improved kernel density estimator for home range

**DOI:** 10.1101/2022.08.21.504698

**Authors:** Jayanth Vyasanakere, Jayanti Ray-Mukherjee, Rahul Joseph Fernandez

## Abstract

1. One of the central challenges in ecology and animal behaviour is to generate animal home range estimations. Kernel density estimate for home range has been one of the most widely used estimates these last few decades, despite its limitations. More recently a network-based kernel density (NKDE) approach has been proposed, which uses Delaunay triangulation.
2. Here we show that NKDE has a discontinuous kernel density. We then develop a new network-based method that emphasises entirely on the edges, instead of the nodes in the network. We call this Edge-Focused Network-based Kernel Density Estimation (EFNKDE). In this method, a unit weight is distributed uniformly along each edge of the network and Euclidean distance is used to compute the contribution of each segment of the edge to the kernel density at a given point.
3. This is a network based method that leads to a continuous and differentiable kernel density. An analytical expression for the same is obtained for the Gaussian kernel. By taking concrete examples and studying different methods and through different perspectives across a range of bandwidths, we show that EFNKDE has many advantages over other methods. We present a model that provides a theoretical basis to the effectiveness of EFNKDE in linear regimes.
4. EFNKDE is easy to apply, can work with minimal data, provides a smooth kernel density that highlights the network and does not overemphasise the data. This method is most suitable for estimating home ranges with narrow corridors and forbidden regions.

## I. INTRODUCTION

Animals’ home range estimation provides critical information of their movement patterns and their habitat preferences that imprints a cognitive map of their home region and the places they would visit (Burt 1943^4^, Powell & Mitchell 2012^18^). Therefore, a home range represents an interplay between the animals’ environment and their understanding of that environment shaped by natural selection to increase an animal’s fitness by optimising space and resource use. For several decades, ecologists have tried to quantify animals’ home range. With the advancements in tracking and telemetry technologies, they have achieved more sophistication in the rate and accuracy of collecting data. This necessitated the progression of new computational techniques that offered elegant insights into animals’ habitat use and home range patterns (Kie et al. 2010^12^, Downs et al. 2011^8^). Critical mathematical and statistical underpinnings have always imposed challenges on home range estimation, where each estimator has been built upon the shortcomings of its predecessors. These methods also extend to the fields of economics, geography and urban planning, and in the fields of computer science and artificial intelligence (Oyang et al. 2005^17^, Timothee et al. 2010^23^).

The earliest method used in estimating home ranges was by using spatial points to construct a minimum convex polygon (MCP) delineated by encompassing all data points (in this case, animal locations) (Mohr 1947^13^). This was followed by other methods, such as kernel density estimation (KDE), that obtain a probability distribution (Silverman 1986^22^, Worton 1989^25^). Here each data point influences the kernel density (which is the probability density) associated with finding the animal at any given point. In such estimators, the bandwidth (denoted here as *h*) which is a smoothing factor or the length scale over which a data point extends its influence, becomes a critical component (Seaman & Powell 1996^19^, Hemson et al. 2005^11^, Signer et al. 2015^21^).

There have been explorations in the use of network based methods on problems where the domain itself is a network, as in hotspots of traffic accidents, street crimes, gas leakages, etc. (Okabe et al. 2009^16^, Xia et al. 2019^26^). Network-based kernel density estimator (which we call NKDE-1) was developed for analysing even two-dimensional planar patterns of data where a network, such as Delaunay triangulation (DT), is drawn taking the data points as nodes and the distance from any given point to different nodes are measured along the network, after reaching the nearest node (Downs & Horner 2007^6^). The network based methods are supposed to bring out the prominence of the network. However, the kernel density obtained is discontinuous. Also, the nodes still get a lot of prominence when *h* is kept small and when it is increased, the features of the network get blurred.

There have been developments which take into account the times at which the data points were captured and effectively handle the correlation effects (Fleming et al. 2015^9^,Noonan et al. 2019^15^). Here we have focused on those methods that take independent and identically distributed data (IID) as input.

In this work we propose and develop a new method that is based directly on the edges of a network, instead of the nodes and produces a continuous and differentiable kernel density. We call it the edge-focused network-based kernel density estimation (or EFNKDE). In Section II, we explain the strengths and weaknesses of some of the kernel density estimators – the original (KDE), more recent (NKDE-1) and its variation that we have worked out (NKDE-2). The formulation of the proposed (EFNKDE) method is detailed in Section III and its few important features are explained in Section IV. We use a readily available online data set to demonstrate the performance of each of the methods - visually in Section V and using a quantitative metric in Section VI. Section VII provides an analytically solvable model to show that the results have a sound theoretical backing. In Section VIII we employ suitable examples to illustrate the usefulness of EFNKDE, even when the domain contains forbidden regions.

## II. A BRIEF OVERVIEW OF OTHER HOME RANGE ESTIMATION METHODS

In order to elaborate on the methods, we have prominently used the two dimensional point data of woodland caribou (Rangifer tarandus caribou) from Northern British Columbia (BC Ministry of Environment 2014^1^, Seip and Price 2019^20^). The original data is huge, consisting of many location tags of many caribous over an extended period of time. From this, we randomly selected a 100 point subset (see Fig. 1(a)) of a single caribou individual assuming that it represents IID. We have then analysed the different home range estimators, highlighting the basic differences across methods and their shortcomings.

**FIG. 1.**
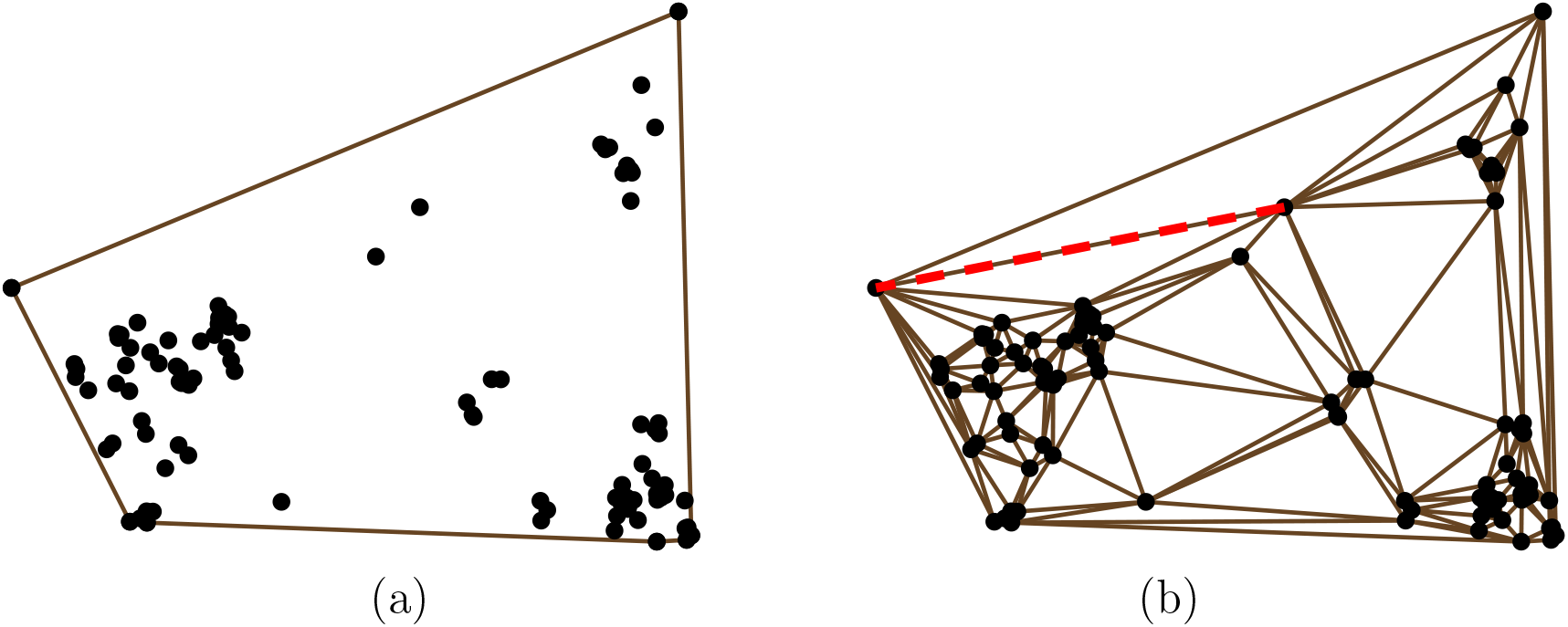
(a) The point data studied in this work along with the minimum convex polygon (MCP) enclosing it. The horizontal and vertical ranges here are 21.1 km and 16.4 km respectively. (b) Delaunay Triangulation (DT) drawn on the same data. The edge highlighted as dashed-red line in (b) will be used in Fig. 5.

The simplest method of home range estimation is to draw MCP that is the smallest polygon where no angle exceeds 180 degrees enclosing all the data points and considering the interior region as the home range (see Fig. 1(a)). Here, no point within the polygon is taken into account and hence MCP is insensitive to any change in the internal region, and thus often considered to be subject to unpredictable bias. Such biases would impose obstacles in understanding important ecological questions, like in understanding how the habitat use patterns change over seasons or between individuals (Nilsen et al. 2008^14^, Burgman & Fox 2003^3^)

A more informative approach is to draw Delaunay triangulation (DT) on the point data (see Fig. 1(b)). DT is a way of triangulating the given set of points in such a way that no point gets inside the circumcircle of any triangle. For the data that we have used, there are 100 nodes, 290 edges and 191 triangles in this DT. There is a larger probability of spotting animals within smaller triangles and the network indicates the possible corridors underlying the data and hence DT is more informative than MCP. This is also the starting point for the network-based methods discussed later in this paper.

The next set of methods estimate the kernel density at *each* point in the region based on the point data, i.e., the home range is described as a probabilistic distribution of animals’ space utilisation. The simplest among them is called KDE (Silverman 1986^22^, Worton 1989^25^). Here, if *d*_*j*_ is the distance from a point where the kernel density is to be obtained, to a point *j* in the point data, then the kernel density at that point is proportional to 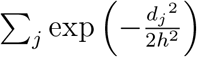 where the Gaussian kernel is used and *h* is the bandwidth. With a small value of *h*, the finer details can be observed, which become obscure with larger values, where only more prominent features are projected. The KDE popularised the literature on home range analysis and is analysis-friendly. However, it has its own limitations (Blundell et al. 2001^2^, Hemson et al. 2005^11^, Downs & Horner 2008^7^). We will later illustrate some of the limitations through specific examples.

Next are network-based methods which draw a network on the point data (like DT in Fig. 1(b)) and build on it. Network-based kernel density estimation (NKDE-1) is a method which is analogous to KDE, but the distances to different nodes from a given point are obtained along the network (Downs & Horner 2007^6^). Here, the nearest node is identified from a given point and from there, the shortest path along the network is taken to any other node. The combined path length plays the role of *d*_*j*_ in the earlier equation. Naturally, this scheme puts the network on the foreground. However, since such a distance is obtained by first locating the nearest node and different nodes can be the nearest ones for two points that are arbitrarily close to each other, the kernel density obtained is discontinuous (see figures 2 and 5).

**FIG. 2.**
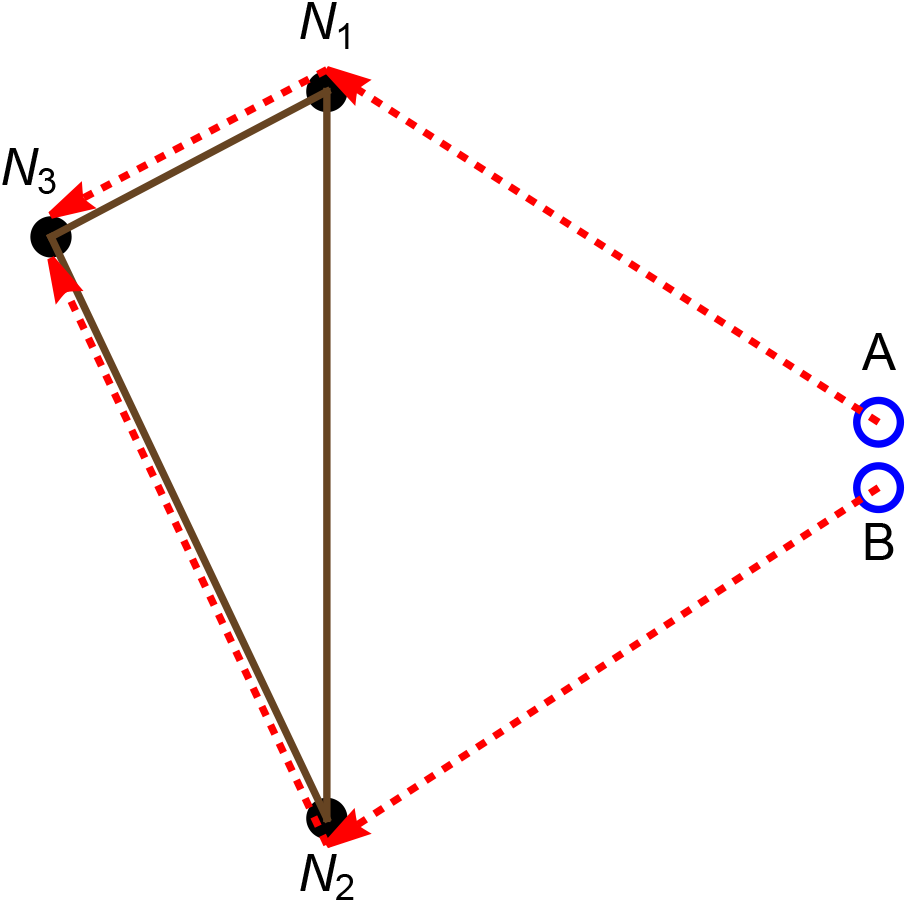
The source of discontinuity in NKDE-1. *A* and *B* are two points at which the kernel densities are to be obtained. *N*_1_ is the closest node from *A* and *N*_2_ is from *B*. Hence the combined path lengths from these points to the node *N*_3_ are *AN*_1_*N*_3_ and *BN*_2_*N*_3_ (indicated by dashed-red lines). These path lengths have a finite difference although *A* and *B* can be arbitrarily close.

We have also developed NKDE-2, which is a slight modification of NKDE-1. Here, from a given point, the nearest point on the network (which may either be a node or a point on an edge) is chosen and the shortest path along the network is taken to any other node. The combined path length plays the role of *d*_*j*_. This also does not give a continuous function because, similar to the argument made earlier, different points on the network can be the nearest ones for two points that are arbitrarily close to each other. Moreover, consider a situation, where the point at which the kernel density is to be evaluated and a node are very close to each other, but an edge in the network intersects the line joining them (see Fig. 3).

**FIG. 3.**
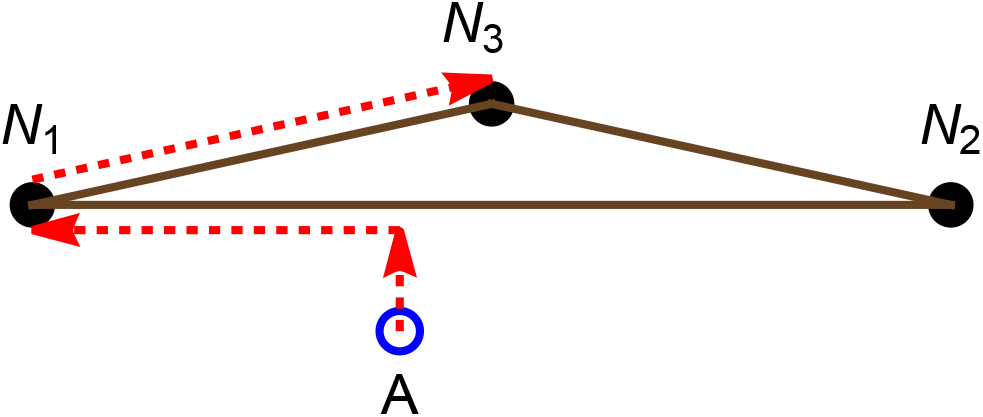
An additional problem with NKDE-2 apart from the discontinuity of the kernel density. *A* is a point at which the kernel density is to be obtained. As per NKDE-2, to reach *N*_3_ from *A*, one has to first reach the nearest point on the network and then find the shortest path along the network. This route is indicated by the dashed-red line. This path length is much larger, compared to the Euclidean distance between *A* and *N*_3_ as one would take in KDE, thereby depleting the kernel density at *A*.

Then, NKDE-2 puts a low kernel density owing to a roundabout route compared to KDE or NKDE-1. So, such points get underrepresented (see the middle part of the dashed-red curve in Fig. 5). In effect, the kernel density gets inflated near the nodes, despite the presence of an underlying network.

Most studies and reviews on this topic highlight MCP and KDE as the popular estimators of animals’ home range. Due to its ease of use, as most GIS tools use KDE, the biases associated with it are often ignored. Also, compared to KDE the network-based methods are rarely used in the field of ecology, due to multiple reasons. First, the network based methods developed so far bring discontinuity along with them as explained above. Secondly, the advantages of using a network has not been convincingly demonstrated so far. Third, implementing network based methods is mathematically more involved than implementing KDE. Here we provide a significant improvement in network-based analysis, where the issues associated with the discontinuities of kernel density have been overcome, the advantages of using the network has been highlighted and the ease of use in EFNKDE is comparable to that in KDE. We have also provided codes to further facilitate the implementation.

## III. FORMULATING EFNKDE METHOD

EFNKDE also starts with the construction of a network like DT over the point data (while the method can be easily extended to other networks or modifications of DT as well, as demonstrated in Section VIII). However, as the name suggests, we do not consider nodes, but focus entirely on the edges of the network. On each given edge, a unit weight is distributed uniformly. Hence the linear weight density on an edge is inversely proportional to its length. We then integrate the weighted contributions coming from each segment of each edge to the kernel density at a point by taking the Euclidean distance between the point and the segments. Here we have used the Gaussian kernel.

The calculation of the kernel density *K* at a point ***r***, denoted as *M* in Fig. 4, which lies on the plane of the point data is as follows. Consider the *i*^th^ edge *E*_*i*_ (with length *L*_*i*_) out of a total of *N* edges in DT. Let *F*_*i*_ be the foot of the perpendicular drawn from *M* to the straight line containing *E*_*i*_ and let *p*_*i*_ be the length *MF*_*i*_. Let *λ* be a coordinate on the line containing *E*_*i*_ such that *λ* = 0 corresponds to *F*_*i*_. On this line, *E*_*i*_ extends from *λ* = *q*_*i*_ (minimum value of *λ*) to *λ* = *q*_*i*_ + *L*_*i*_ (maximum value of *λ*). Note that *p*_*i*_ is a non-negative real number, while *q*_*i*_ and *q*_*i*_ + *L*_*i*_ can be any real numbers. Both *p*_*i*_ and *q*_*i*_ depend on *E*_*i*_ and *M* being considered.

**FIG. 4.**
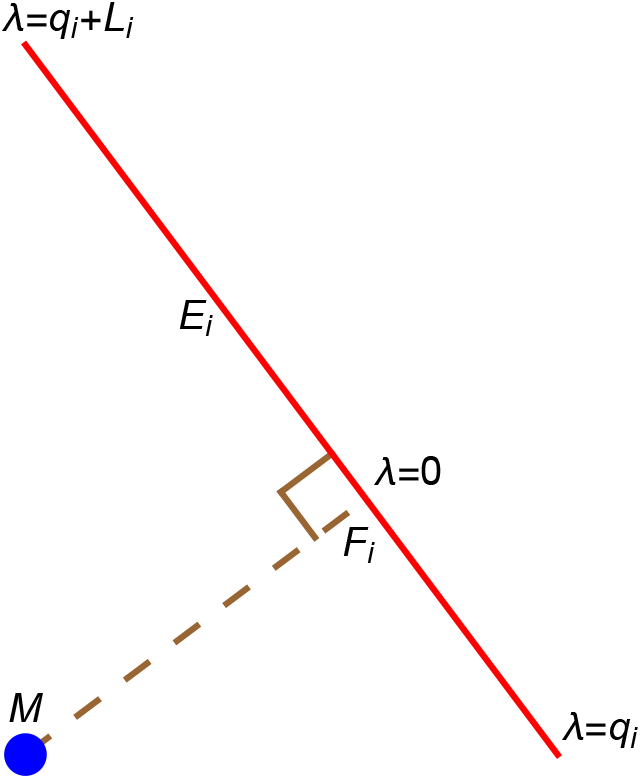
Notations used in EFNKDE leading to equations 1 and 2. See text for explanation.

The weight in an element *δλ* which is a part of *E*_*i*_ is *δλ/L*_*i*_. Given the Gaussian kernel with band width *h*, the contribution *δK* of this element to *K*(***r***) is proportional to

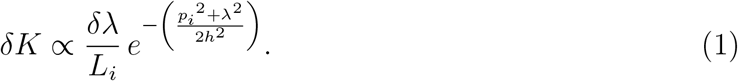

For the full contribution of *E*_*i*_ to *K*(***r***), this has to be integrated over *λ* ranging from *q*_*i*_ to *q*_*i*_ + *L*_*i*_, which gives rise to the error function (erf). Thus, by including all the edges and along with the normalisation (which means that the integral of *K*(***r***) over the plane is unity),

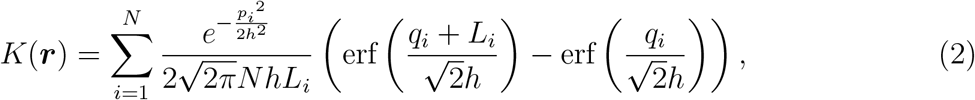

where the dependence on ***r*** comes through *p*_*i*_ and *q*_*i*_ (see appendix A). We thus have an *analytical* expression for *K*(***r***) which is bounded, positive, continuous, differentiable and normalised. The continuity and differentiability of *K*(***r***) follows from those of the exponential and error functions.

The contribution of each *E*_*i*_ decays exponentially with *p*_*i*_. The derivative 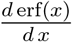 decreases monotonically with the absolute value of *x*. This ensures that the contribution of *E*_*i*_ to *K*(***r***) increases with the proximity of its centre to *F*_*i*_. As *L*_*i*_ *→ ∞*, the term within the brackets in Eq. (2) increases and saturates, but the overall contribution of *E*_*i*_ *→* 0 because of the dilution of weight density (*L*_*i*_ present in the denominator). All these are intuitive and desired properties of a kernel density function.

## IV. A FEW IMPORTANT FEATURES OF EFNKDE

Here we discuss a few elegant features implicit in Eq. (2). The first point to note is that the Kernel density obtained using EFNKDE gives emphasis on edges instead of nodes. This feature of EFNKDE leads to several benefits in the applications as discussed in later sections. As pointed out earlier, the other feature is that the kernel density is smooth (continuous and differentiable). These two features are illustrated in Fig. 5, where the variation of kernel density along an edge in DT (the one highlighted in Fig. 1(b)) is shown. Except for EFNKDE, the nodes (*ξ* = 0 and *ξ* = 1) are prominent in all the other methods. The discontinuity in NKDE-1 can be seen as one approaches the nodes other than the corners of the edge highlighted. The reasons for these behaviours were discussed earlier in Sec. II.

**FIG. 5.**
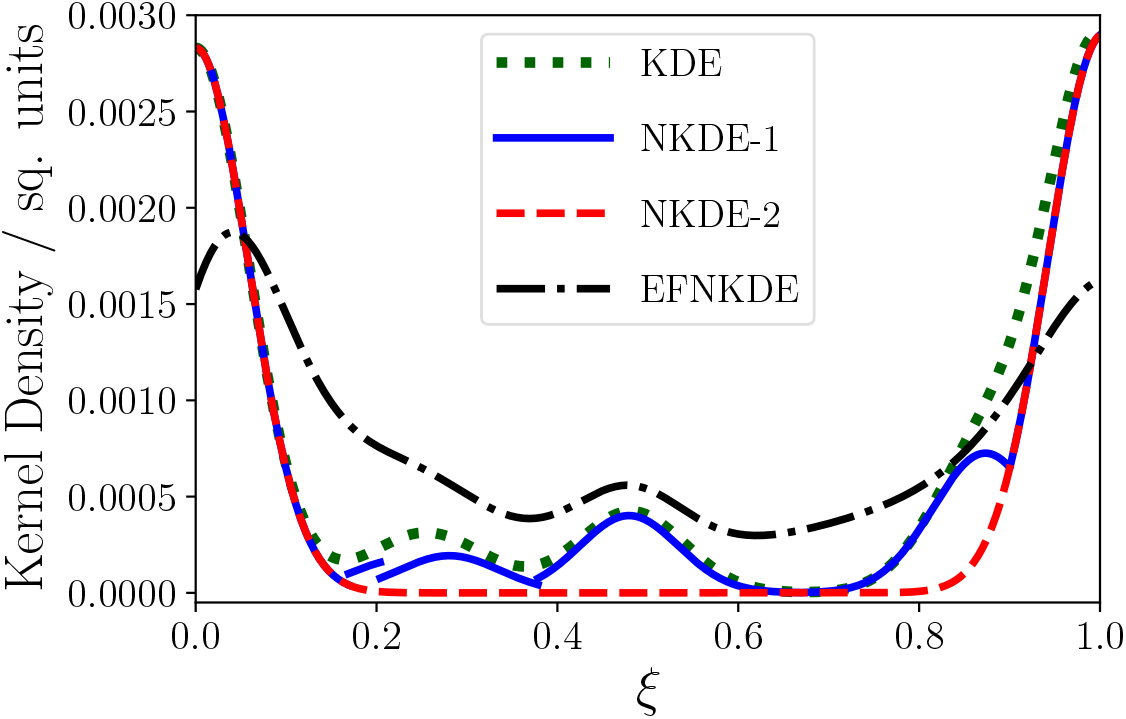
Kernel density along an edge in DT as predicted by different methods. The variable *ξ* takes values from 0 to 1 as one moves from the left corner to the right corner on the edge highlighted in Fig. 1(b). Bandwidth *h* is kept the same (0.75 km) for all the curves. As expected, the curve corresponding to NKDE-1 is discontinuous. It can be seen here that EFNKDE does not give too much prominence to nodes (*ξ* = 0 and *ξ* = 1).

The third advantage of EFNKDE is that while it inherits the primary feature of the other network-based methods, which is to highlight the network, it is more analysis-friendly compared to them, as much as KDE. This is because the Euclidean distances from the segments of the edges are used, instead of taking the distances along the network. For example, the normalisation could be performed analytically in Eq. (2) for EFNKDE. Whereas, for the other network-based methods it is not possible to obtain such closed form expressions.

## V. APPLICATION OF DIFFERENT METHODS TO REAL DATA: VISUAL COMPARISON

We now discuss the performances of different methods from different perspectives. To begin with, in Fig. 6, we apply the different kernel density methods to the data shown in Fig. 1 and visually compare them. While the kernel densities produced by both NKDE-1 and NKDE-2 are discontinuous, that in NKDE-1 is pronounced. One can see from the last row that in EFNKDE, the kernel density is continuous and differentiable and the nodes do not show up prominently even for smaller bandwidths, thereby allowing the network to get highlighted. Consequently, the points that are not close to a node also get a significant density, still retaining the details of the underlying pattern. This also means that the density persists, even if barely, in regions with low occurrence of points. This is beneficial because this method can capture potential space use patterns even with a minimal sample of points. We can see this in Fig. 6, especially exemplified in the lowest bandwidth configurations, when we compare EFNKDE to other methods. Longer edges getting lower weights is also a desirable property.

**FIG. 6.**
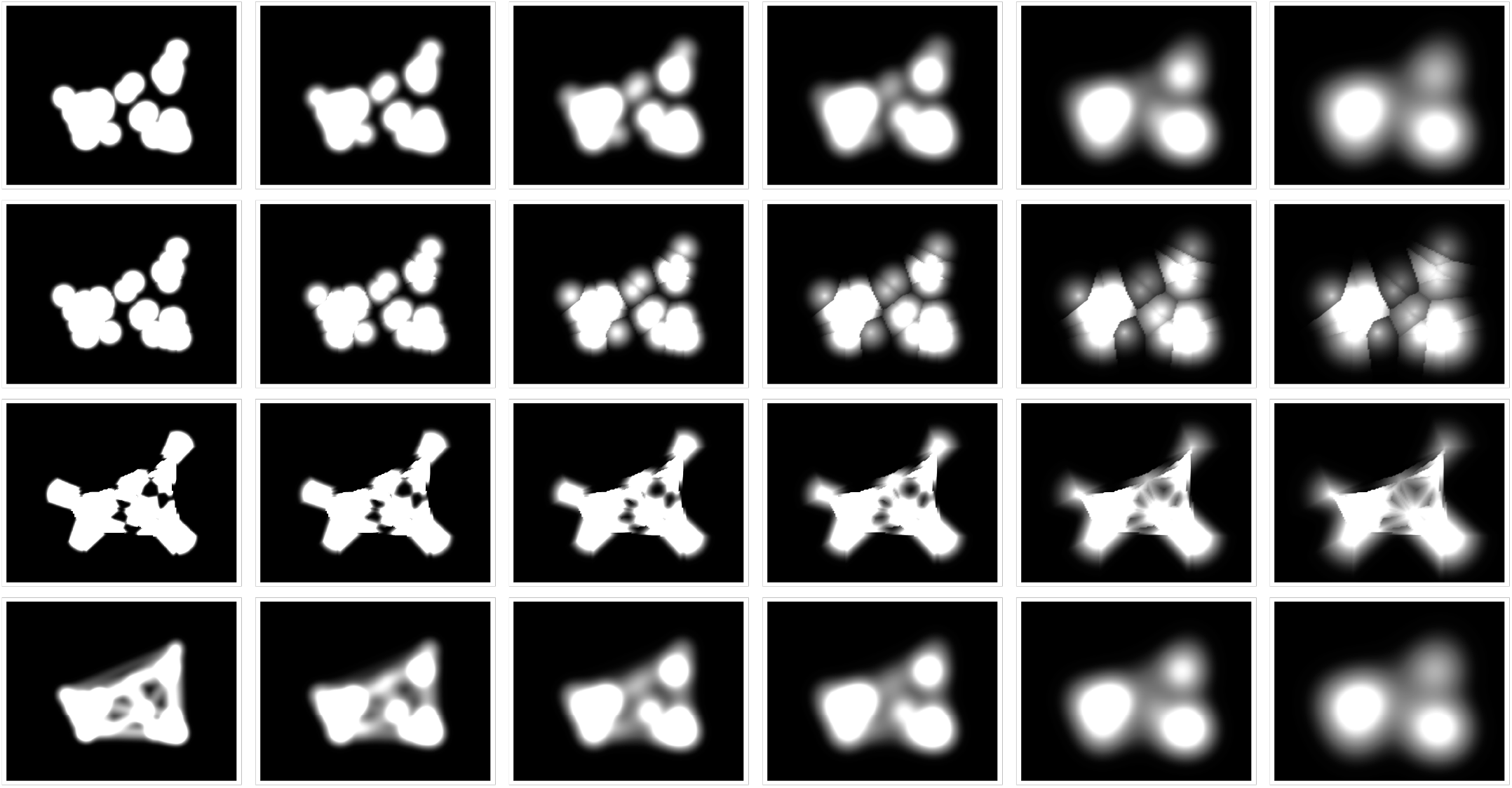
Comparison of kernel densities obtained from different methods for varying bandwidths. The top row corresponds to KDE, the second row to NKDE-1, the third row to NKDE-2 and the bottom row to EFNKDE. In each row, the bandwidth *h* increases from a small value (left) to a large value (right); the precise values (in km) for different columns being 0.7, 1.0, 1.3, 1.6, 2.2 and 2.8.

## VI. APPLICATION OF DIFFERENT METHODS TO REAL DATA: QUANTITATIVE COMPARISON

As noted earlier, EFNKDE promises to be useful when the data is sparse. We demonstrate below that it is most useful when the distribution is linear, which is also the regime where node-based methods like KDE may not give satisfactory results. For this we continue to work with the same caribou data (Fig. 1(a)), but we compress the distribution along the *y* direction (vertical direction), keeping it intact along the *x* direction (horizontal direction). Parameter *µ* is the factor by which the *y* range gets multiplied. This means, *µ* = 1 is the original distribution. As *µ* is reduced, the distribution shrinks along the *y* direction to become more and more linear. We show below that EFNKDE outperforms the other methods in this regime.

We take 25 random points (called training set) from the data (which totally has 100 points) and a given method is employed to obtain the kernel density. The result is then tested on the remaining 75 points (called the test set). We construct 50% isopleth (which means the smallest region having 50% probability under the kernel density obtained). We call the percentage of points from the test set falling within this region as *P*_50_. In a similar way, we construct 95% isopleth and obtain *P*_95_. This method is called split-sample crossvalidation. We have chosen 25 and 75 points in the training and the test sets to make the training data more sparse (where EFNKDE is most promising) and the test criteria finer.

Ideally *P*_50_ should be 50 and *P*_95_ should be 95. Before we check how different methods perform, we obtain the optimum bandwidth using the likelihood cross-validation method (Silverman 1986^22^). In the case of KDE, NKDE-1 and NKDE-2, we omit one of the data points from the training set and construct the respective kernel density. We can evaluate this kernel density at the location of the omitted data point to obtain the likelihood associated with that point. The optimum bandwidth is the one that maximises the product of such likelihoods taken over all the points in the training set. This algorithm gives exactly the same bandwidth for both NKDE-1 and NKDE-2. In the case of EFNKDE, we follow the same algorithm, except that for each data point, we omit all the *edges* containing that point to evaluate its likelihood.

Using this algorithm, the bandwidths were obtained for different values of *µ* (Fig. 7(a)).

**FIG. 7.**
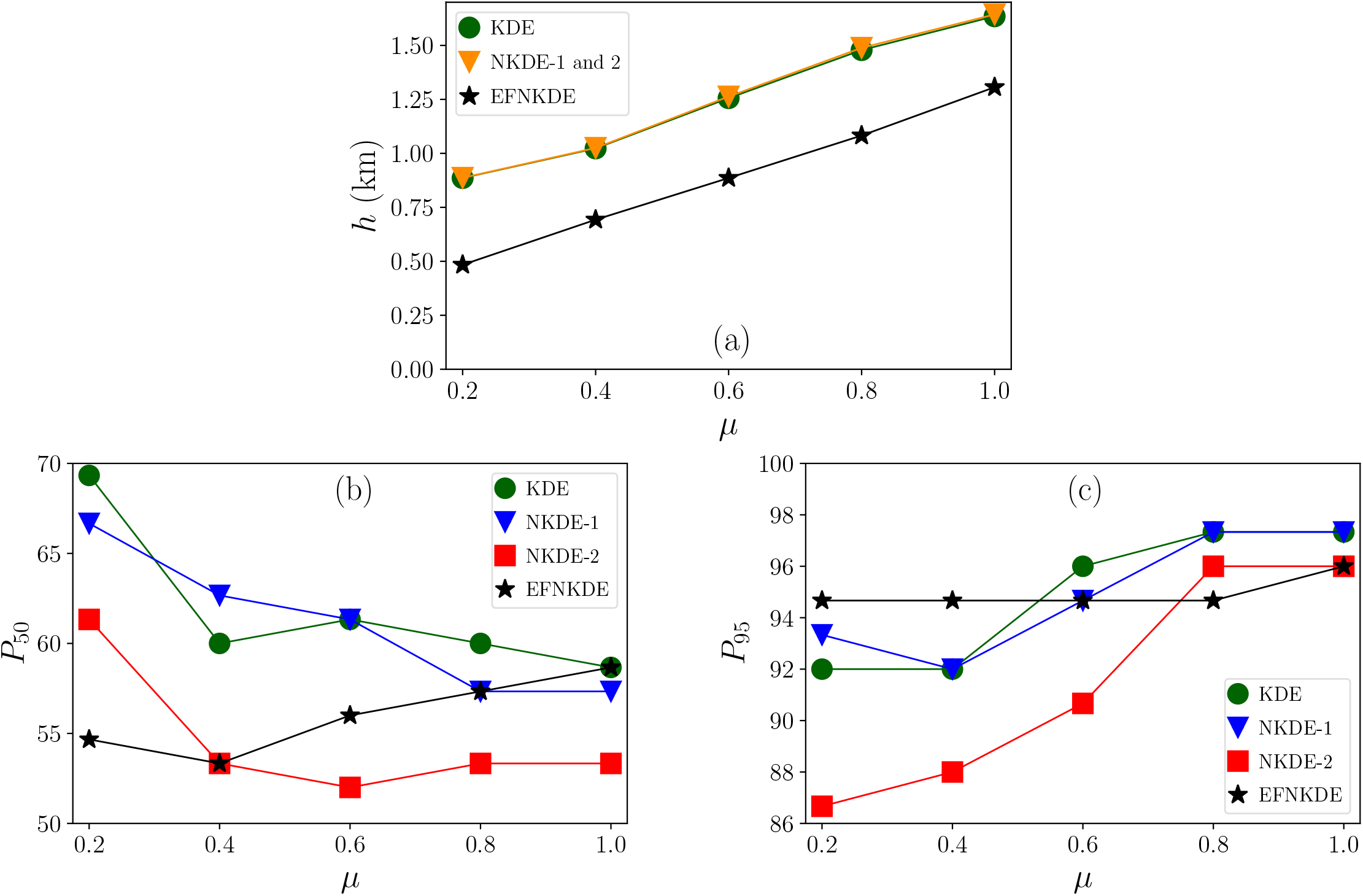
Performance of different methods when *µ* (the factor by which the caribou data is multiplied along the *y* direction) is varied. (a) Optimum bandwidth obtained from likelihood cross-validation method. (b) and (c) show the percentage of points from the test set falling within 50% and 95% isopleths respectively. For smaller *µ* (more linear distribution), EFNKDE has *P*_50_ close to 50 and *P*_95_ close to 95.

With that, the actual values of *P*_50_ and *P*_95_ were obtained for different methods (Fig. 7(b) and 7(c)). It is apparent that for smaller *µ*, it is EFNKDE whose *P*_50_ is close to 50 while simultaneously having *P*_95_ close to 95.

Why is it that EFNKDE continues to perform well at small *µ*, while the node-based methods struggle? In Fig. 8, the isopleths along with the distribution of data points are shown for a fixed small *µ*. Note that the distribution is not similar along the *x* and *y* directions. Let us consider KDE for example. A small bandwidth would miss out points along the *x* direction, while a large bandwidth will unnecessarily extend the home range along the *y* direction. Similar statement is true for NKDE-1 and NKDE-2 also. Therefore these methods face conflict and will have to go for a compromise. On the other hand, for EFNKDE the advantage comes from the fact that it is network based and more importantly edge focused. The network extends largely along the *x* direction and can get closer to all the points. Therefore with a smaller bandwidth it can capture the distribution along the *y* direction. That EFNKDE operates with a smaller bandwidth compared to other methods in such a situation is also evident from Fig. 7(a).

**FIG. 8.**
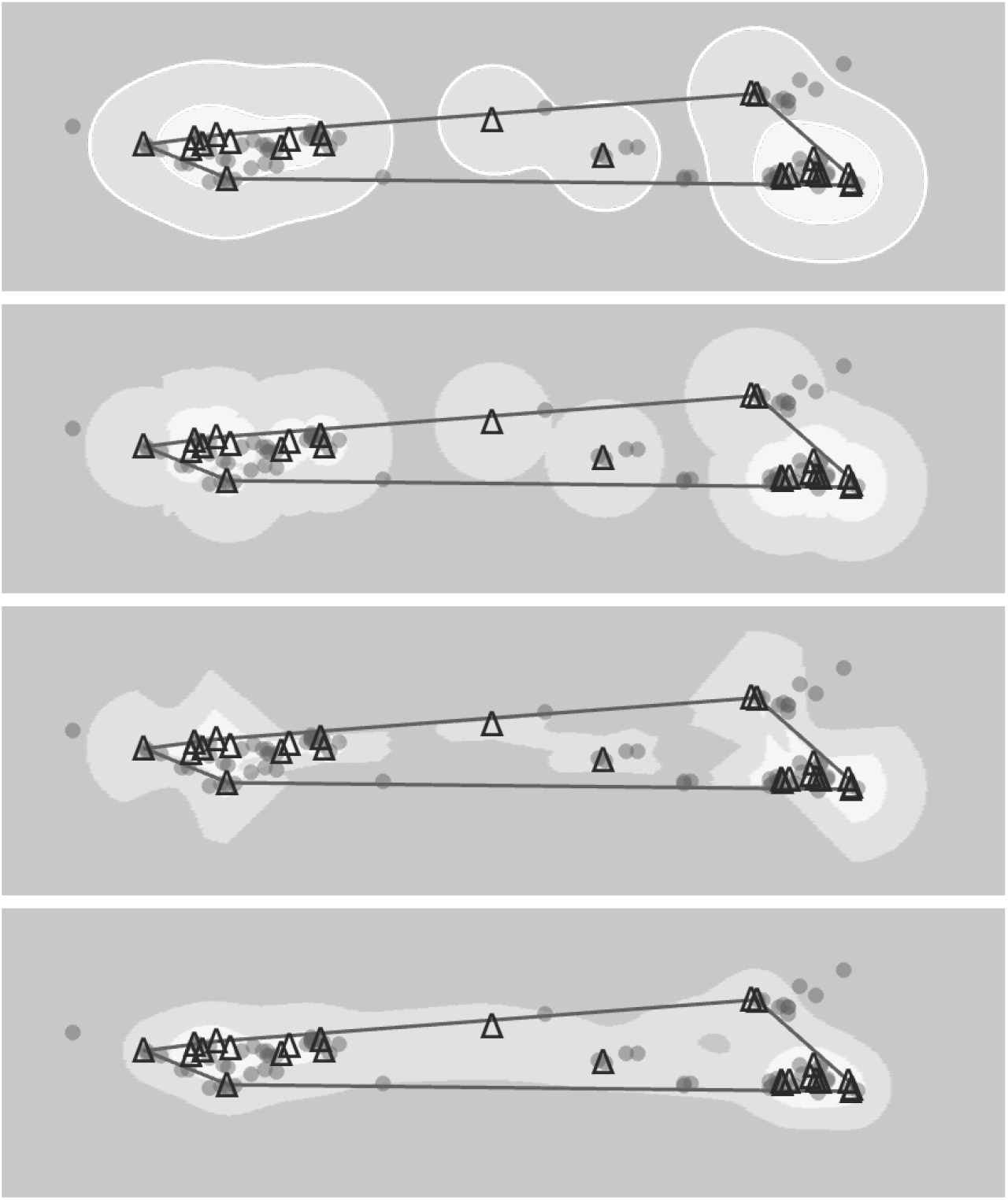
Figures showing 50% and 95% isopleths for *µ* = 0.2. The darkest region is out of 95% isopleth. The lightest region is within 50% isopleth. The region with intermediate darkness is outside 50% isopleth but inside 95% isopleth. The triangles represent the data points from the training set and circles represent those from the test set. MCP constructed from the training set is also shown. The top row corresponds to KDE, the second row to NKDE-1, the third row to NKDE-2 and the bottom row to EFNKDE. In each case the bandwidth employed is from the likelihood cross-validation method as shown in Fig. 7(a). Here it can be seen that the node-based methods, unlike EFNKDE, overestimate the home range along the *y* direction.

The success of EFNKDE is not an artifact of the specific bandwidth-selection algorithm (likelihood cross-validation) used. To prove this, one could in principle consider different methods to calculate bandwidth. However, the different methods yield only specific values for the bandwidth. Since there is no end to the list of methods to calculate the bandwidth, checking the results by employing each of them is both tedious and insufficient. Hence we adopt the following strategy, which serves the stated purpose.

When the variation of *P*_50_ and *P*_95_ with bandwidth is considered (Fig. 9) for a fixed value of *µ* = 0.2, it can be seen that irrespective of the bandwidth chosen, node-based methods cannot outperform EFNKDE, i.e., they struggle to simultaneously maintain *P*_50_ at 50 and *P*_95_ at 95. For these methods, a smaller bandwidth imposes a drop of *P*_95_ and a larger bandwidth would impose a rise of *P*_50_.

**FIG. 9.**
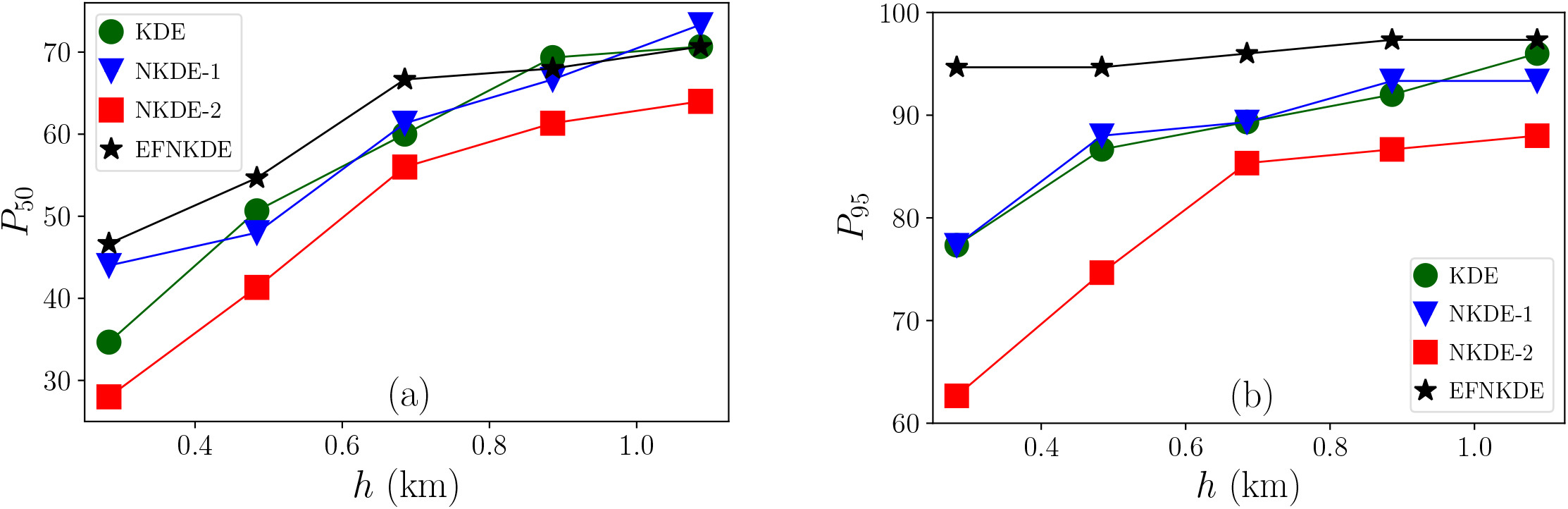
Variation of *P*_50_ and *P*_95_ with bandwidth for *µ* = 0.2 in different methods. (a) For higher bandwidths, *P*_50_ tends to rise well above 50. (b) For lower bandwidths, *P*_95_ tends to fall below 95. EFNKDE finds the best balance.

## VII. QUANTITATIVE COMPARISON: A TOY MODEL WITH AN INFINITE LINEAR CORRIDOR

To illustrate the arguments of the previous section for a linear distribution even more quantitatively, motivated by Fig. 8 we further build the following toy model, where the true density is a-priori known. This model captures the essential feature (linearity) of the distribution in Fig. 8 that is giving EFNKDE an advantage and considers a simpler situation showcasing that feature. The purpose is to gain a strong control over analysis, thereby establishing the benefits of EFNKDE beyond any doubt, including those related to bandwidth selection.

Imagine an infinite linear corridor shown as a shaded region in Fig. 10. We call the direction along the corridor as *x* and that perpendicular to it as *y*. We are given with the data points that are located at (*ai*, 0), where *i* runs over all the integers. Thus the points are uniformly spaced with spacing *a*. The width of the corridor is *b*. DT is obtained by joining all the data points as shown. We also have the luxury of knowing the true density that is given by

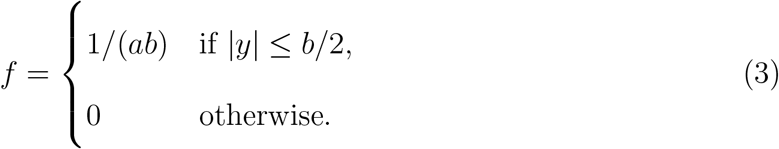

**FIG. 10.**
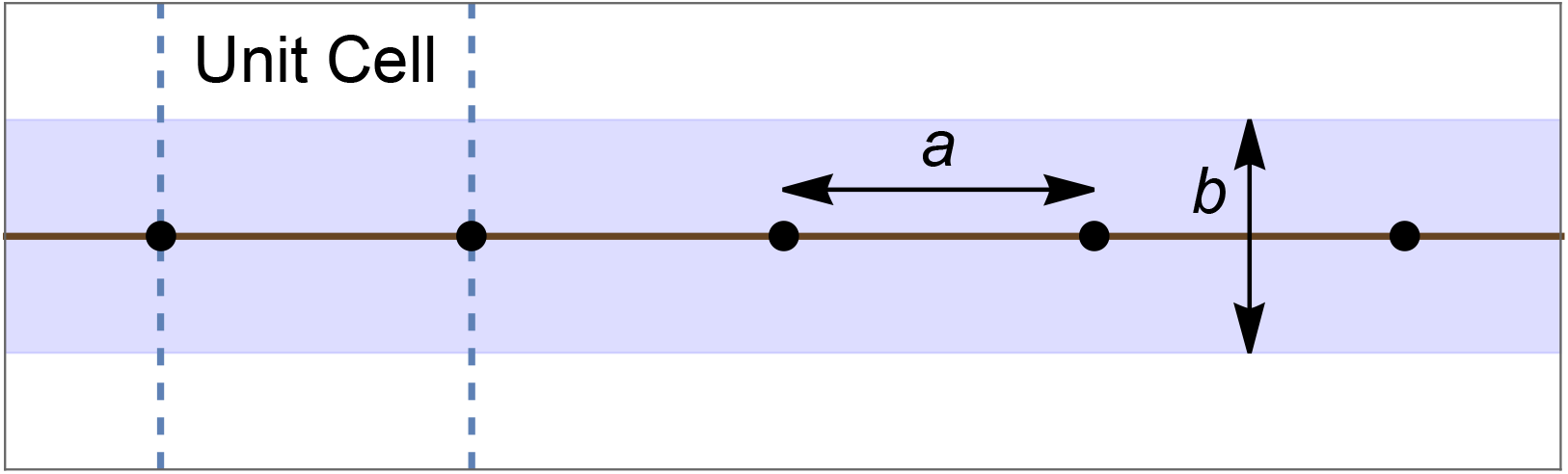
An infinite linear corridor with uniform density within the shaded strip that extends from *x* = *−∞* to *x* = *∞*. The data points (black dots) are uniformly spaced, with spacing between the successive points as *a*, and the width of the corridor is *b*. The solid horizontal lines connecting the dots represent DT. The region between the two vertical dotted lines represents a unit cell.

The question now is how well do different methods perform in reproducing the true density based on the data given?

Before applying different methods, three technical points are worth mentioning. First, it is important to note that the normalisation will have to happen per unit cell, which is the region between two lines parallel to the *y* axis and separated by a distance *a*. A naïve application of the formulae (which is normalised to the entire *x − y* plane) would yield zero as the kernel density owing to the total number of data points being infinite. Second, scaling the value of the true density within the strip by a constant is not going to have any impact on any of the results presented here. Third, if one is wondering where the *triangles* are in DT, one can think of an extra single point at *y* = *∞*. This point connects to every data point lying on the *x* axis through lines that are parallel to the *y* axis. These form triangles. Since the length of the sides parallel to the *y* axis would be infinite, they will not contribute to the kernel density and hence one need not worry about them.

Let *f*_*h*_ be the density function obtained through a given method for a given value of bandwidth *h*. In the case of KDE, the sum over the data points can be done analytically to yield

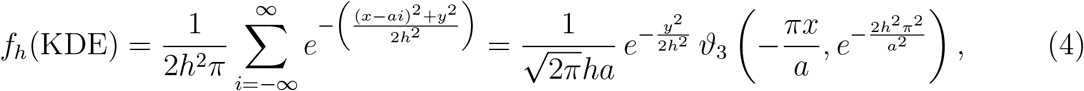

where *ϑ* is the elliptic theta function. In the case of EFNKDE, referring to Eq. (2), *p*_*i*_^2^ = *y*^2^ for all *i* and the contribution from the neighbouring edges will involve cancellations because *q*_*i*_ + *L*_*i*_ = *q*_*i*+1_. Noting that erf(*±∞*) = *±*1, we can again obtain an analytical expression for the density as

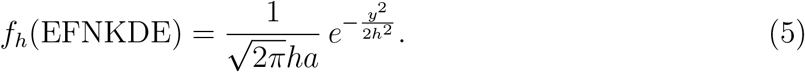

To evaluate the performance, we define a dimensionless measure of root mean squared deviation/error of *f*_*h*_ from *f* as

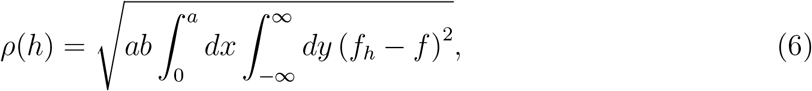

where the integration extends over a unit cell.

The optimum *h* is the one that minimises *ρ*(*h*). For different values of *b/a*, the optimum *h* is plotted in Fig. 11(a). Consistent with our earlier observation, EFNKDE chooses a lower bandwidth compared to KDE. The minimum value of the error itself is plotted in Fig. 11(b). It can be clearly seen that EFNKDE performs uniformly for all *b/a*, while the error in KDE shoots up as *b/a* tends to 0. Connecting back to applications, this model explains why in the examples considered in Fig. 7, EFNKDE performed consistently throughout, whereas the other methods struggled in the linear regime.

**FIG. 11.**
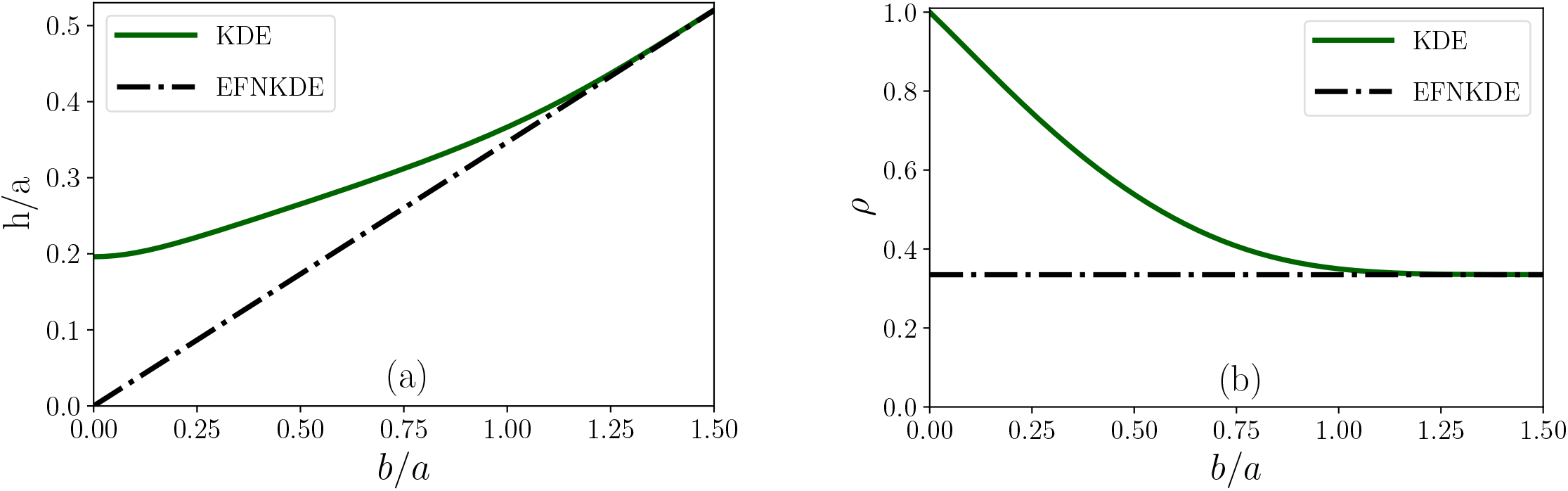
(a) The optimum bandwidth that minimises *ρ* for both KDE and EFNKDE in the infinite linear corridor. In the latter case, *h* = 0.3467 *b* and goes to 0 as *b* goes to 0. (b) The minimum value of the dimensionless root mean squared error for KDE and EFNKDE. In the former case *ρ →* 1 as *b/a →* 0. In the latter case, the error is a constant and is equal to 0.3349. In both the graphs, KDE and EFNKDE results overlap for large *b/a*. EFNKDE can be seen to have a lower error than KDE and this is more pronounced in the small *b/a* regime.

The reason for the difference in the performances is similar to the one that is explained earlier. At small *b/a* KDE cannot achieve uniformity along the *x* direction and simultaneously reproduce the variation along the *y* direction. On the other hand, in EFNKDE the uniformity along *x* comes naturally from the nature of the kernel allowing *h* to decrease arbitrarily to handle the narrow distribution along *y*. These are visible in Fig. 12 even at a relatively large value of *b/a*.

**FIG. 12.**
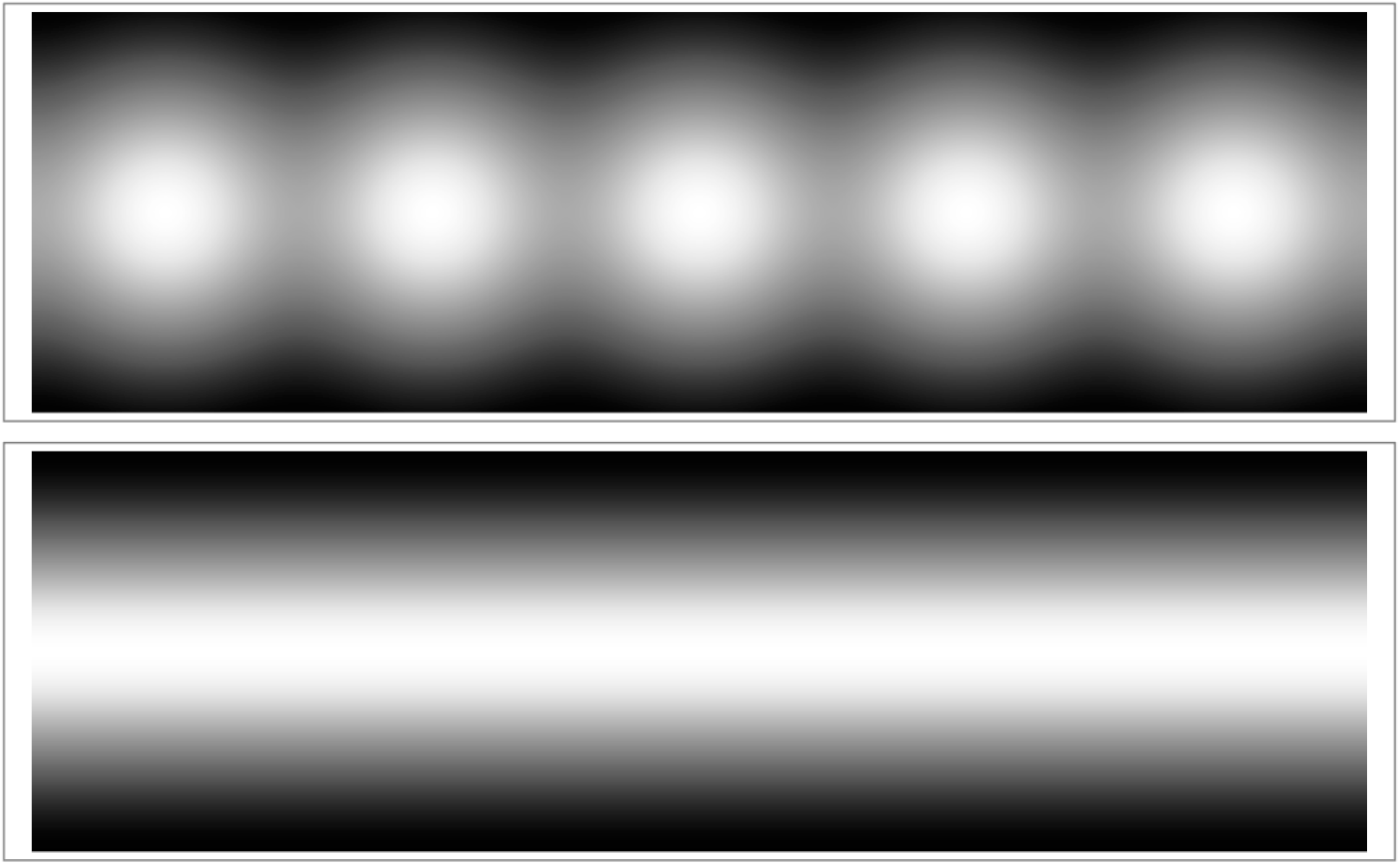
Density plots for *b* = 0.75 *a* in the infinite linear corridor. The top row corresponds to KDE and the bottom row to EFNKDE.

If the other methods that focus on the nodes can be compared with a point source of light (ex: an incandescent bulb or a lamp), EFNKDE being edge-focused can be compared with a linear source of light (ex: a tube light). The results here are equivalent to saying that it is better to light a narrow corridor with a linear source of light than using point sources.

Why do KDE results overlap with EFNKDE at large *b/a*? With a large enough *h* in KDE, the uniformity along the *x* direction is taken care of, irrespective of the precise value of *h*. This removes the necessity to go for a compromise (as explained earlier) and a suitable (large) *h* can be chosen to fit the distribution along the *y* direction. This overlaps with the mandate of EFNKDE in this problem, which explains the overlap in the results at large *b/a*.

Why does *ρ* → 1 as *b/a* → 0 in KDE? Since in this limit, *f* tends to a Dirac delta function along the *y* direction (see Eq. (3)), the dominant contribution to *ρ* comes from the region of the unit cell within the corridor, whose leading term is 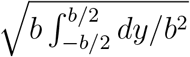, which is 1.

Here we have considered only KDE and EFNKDE, where analytical expressions for the kernel density can be obtained, including the normalisation (see Section IV). It is easy to see that in the small *b/a* regime (which is the important regime for this illustration), NKDE-1 and NKDE-2 are identical with KDE.

One can argue that a node based method like KDE can also be made to perform well in the above situation by having two bandwidths, one for the *x* direction and the other for *y*. However, that strategy cannot work in a more realistic situation where the home range can involve curves that are not necessarily straight lines, as seen in the following examples, where EFNKDE continues to perform well.

## VIII. ABILITY OF EFNKDE TO HANDLE FORBIDDEN REGIONS

One of the major issues with any home range estimation is that the predicted home ranges often extend into forbidden regions, like that of fish extending on to the land for example. EFNKDE (applied as it is) is no exception. Two data points lying far apart may form an edge in DT that passes through a forbidden region. However, this drawback can be easily taken care of in EFNKDE with the following modification.

While the data points (which cannot lie in the forbidden region) come from observations, DT is entirely our construct. So after constructing DT, check visually and knock off all the edges that pass through the forbidden region. With the remaining edges apply EFNKDE as in Eq. (2). This idea is conceptually similar to constructing Urquhart graphs (Urquhart 1980^24^), where the longest edge from each triangle in DT is removed. With this, the drawback mentioned above for EFNKDE can actually be turned into an advantage. To illustrate this, we give two examples, where the data points are plotted and visualised against the background domain consisting of forbidden regions. In both of these cases, the data sets and the boundaries have been simulated to best illustrate the concept.

The first example is that of a lake with spiral arms. Fig. 13(a) shows the data points alone that correspond to crocodiles which are localised to the lake. Fig. 13(b) shows the same along with the lake boundary. In Fig. 13(c), we have constructed DT and in Fig. 13(d) we have knocked off all the edges which go out of the lake (for example those edges joining points from neighbouring spiral arms). With the remaining edges, we apply EFNKDE and the other methods with a suitably chosen bandwidth to obtain the kernel densities as shown in the bottom row of Fig. 13. Remarkably, even in such a challenging situation, EFNKDE has managed to capture the spiral arms of the lake! In the case of KDE, there is no network in the first place. NKDE-1 and NKDE-2, although are network based, have failed to take advantage of the removal of forbidden edges as they are node-focused. The only effect of varying the bandwidth in the other methods is to vary the size of the spots and hence they cannot still reproduce the spiral arms, irrespective of the bandwidth used. Clearly, being edge-focused has given EFNKDE an ‘edge’ over other node based methods.

**FIG. 13.**
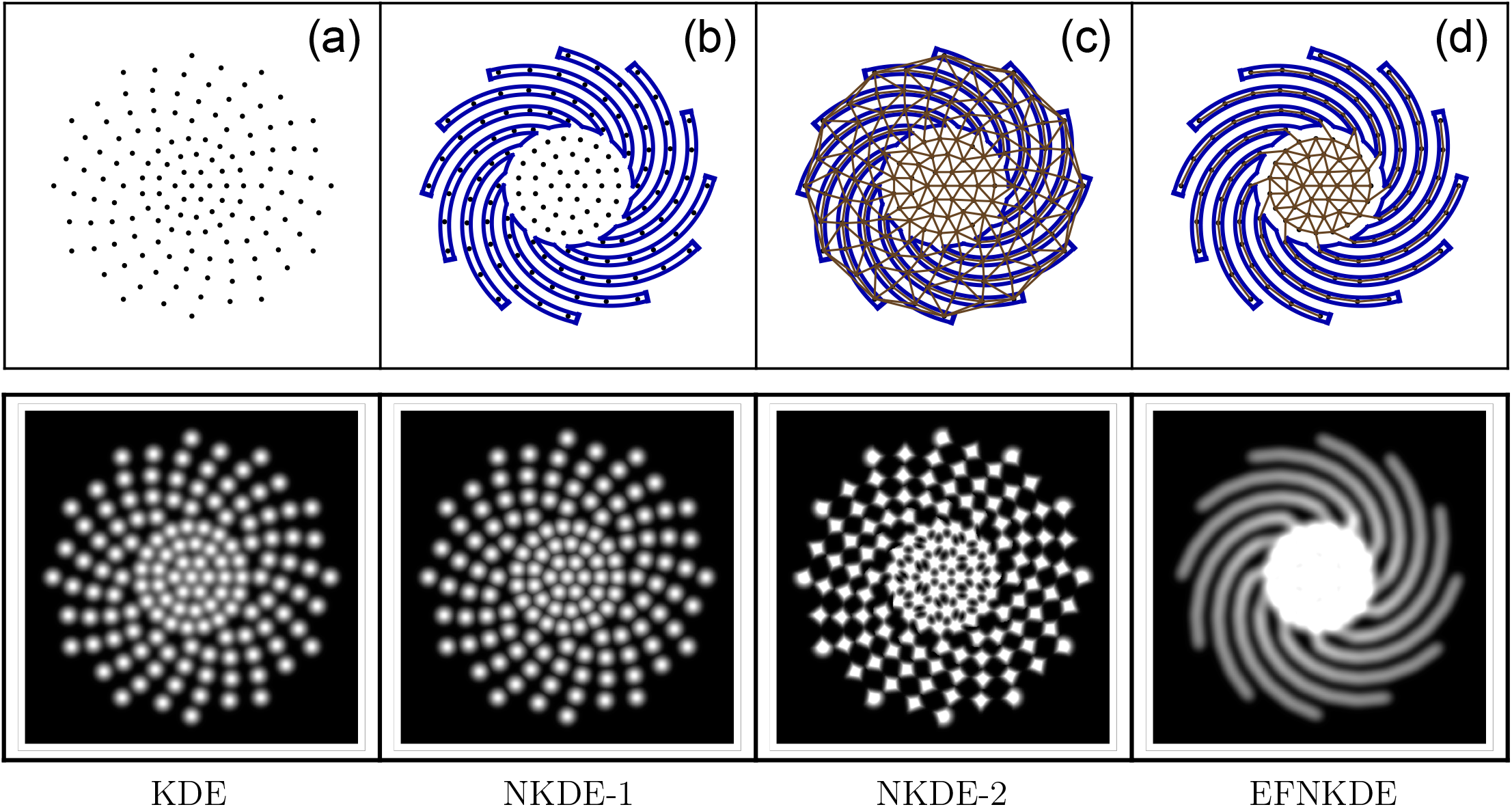
A lake with spiral arms. (a) Data points. (b) Data points along with the lake boundary. (c) DT, with all the edges intact. (d) DT, with edges passing through the forbidden region removed. The resulting kernel densities obtained from KDE, NKDE-1, NKDE-2 and EFNKDE, with the same bandwidth, are shown in the second row. Only EFNKDE could reproduce the spiral structure.

Why can’t KDE be employed in this situation and later make the kernel density zero by hand in the forbidden region? There are two reasons why this is not efficient. First, knocking off edges from EFNKDE can be visually done unlike setting hard boundaries in KDE, where one needs to know the precise functional form of the boundary. Second, within its exponential tail, EFNKDE can in principle capture small excursions into the forbidden region (say crocodiles getting on to the shore). There is no natural way in which setting hard boundaries in KDE can accommodate this.

Encouraged by this success of EFNKDE, we take up the second example where the home range is a union of multiple disconnected regions. The top row of Fig. 14 shows fish as data points in a disconnected lake. The middle row shows DT with edges retained (solid lines) and edges removed (dashed lines). The removed edges are those which have gone out of the lake. With the retained edges, EFNKDE is applied and the kernel density obtained is shown in the last row. Even here EFNKDE has distinctly managed to capture the disconnectedness in the home range. Again, it is easy to see that no node based method could have naturally achieved it.

**FIG. 14.**
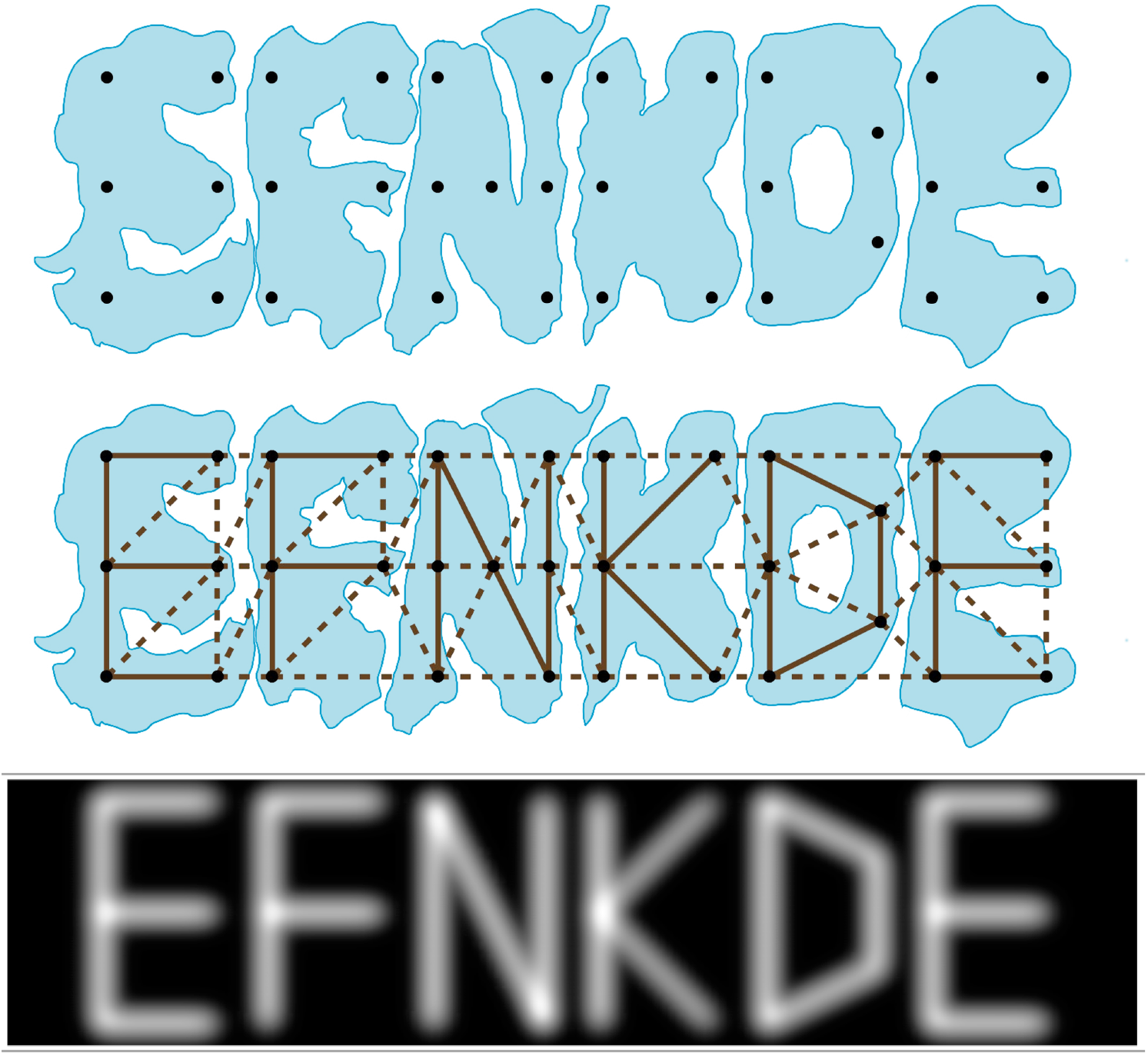
A lake with six disconnected regions. The top row shows the data points along with the boundaries. The middle row also includes DT, with edges passing through the forbidden region removed. Solid lines are the edges retained and the dashed lines are the edges removed. The last row is the kernel density obtained from EFNKDE.

Can one automate the knocking off of the forbidden edges? A more fundamental question to ask here is: “how well-defined is the forbidden region itself?” In a given situation, if there is a code to tell whether a given point in the plane falls within the forbidden region or the allowed region, then from there it is straightforward to automate knocking off of the forbidden edges. In Fig. 13, we knocked off the forbidden edges through a code, because the mathematical equation of the boundary could be obtained. On the other hand, Fig. 14 represents a more realistic example where such a mathematical luxury may not be affordable. There we have removed the forbidden edges manually. The crucial point here is that the user can knock off the forbidden edges *even visually* without getting bogged down by the analytical geometry of the boundary shape. If the boundary is complex, an optimal strategy would be to identify regular regions (like circles or rectangles) that lie completely within the forbidden region and remove the edges passing through these subregions using a code; the remaining forbidden edges can be visually knocked off later.

## IX. CONCLUSION

As suggested by several authors previously, the reflection of a home range of an animal is more complicated than a few pattern assemblages. An animals’ cognitive and memory maps along with their habitat choice and foraging decisions make a home range more complex than it is often projected. Hence the contribution of an expert’s eye and a thorough knowledge of the habitat is undeniable for its practical purposes. Nonetheless, it becomes pertinent to delineate a statistically and computationally sound home range to estimate and compare an animal’s space use across habitat and seasons.

Here, in summary, we have developed a method of home-range analysis (EFNKDE) that focuses on the edges in the network instead of nodes. This is an improvement because the kernel density function obtained is continuous, differentiable (smooth), no region on the plane gets under-represented, the network gets prominence, and the analysis is easy and intuitive. These virtues make it suitable for analysing home ranges with narrow corridors and forbidden regions, particularly when working with minimal set of points.

While formulating EFNKDE, we have given each edge a weight that is inversely proportional to its length. It may be interesting to explore other functions of the length. That would provide an additional knob to tune along with the bandwidth. Here we have not considered methods that include auto correlation (Fleming et al. 2015^9^,Noonan et al. 2019^15^) and methods like local convex hull (Getz and Wilmers, 2004^10^, Chirima and Smith 2017^5^) that are not kernel based. Incorporating these along with EFNKDE in home range estimation can provide scope for further development.

## X. ACKNOWLEDGEMENTS

JV acknowledges generous support from Azim Premji University through a research grant with project ID: Univ RC00266.

## XI. CONFLICT OF INTEREST STATEMENT

There are no conflicts of interest.

## XII. AUTHORS’ CONTRIBUTIONS

JV and JRM conceptualised the idea. JV developed the method, did the calculations, wrote the codes in Wolfram Mathematica and generated the plots. RJF acquired the caribou data, reproduced the codes in Python and R, and generated respective repositories. JV and JRM wrote the manuscript. All authors critically proofread the draft and gave final approval for publication.

## Appendix A: Obtaining *p*_*i*_ and *q*_*i*_

The tricky part in Eq. (2) is to obtain *p*_*i*_ and *q*_*i*_ which are defined in Fig. 4 and the text around it. Let ***r*** be the position vector of a point *M* where the kernel density is to be evaluated and *E*_*i*_ be an edge in DT. Let ***a*** and ***b*** be the position vectors of the corners of *E*_*i*_. Now define the following three vectors: ***v***_*ra*_ = ***r*** *−* ***a, v***_*rb*_ = ***r*** *−* ***b, v***_*ba*_ = ***b*** *−* ***a*** and the following two dot products: *α* = ***v***_*ra*_ *·* ***v***_*ba*_, *β* = *−****v***_*rb*_ *·* ***v***_*ba*_. Then the position vector of the foot of the perpendicular (*F*_*i*_) from *M* to *E*_*i*_ is 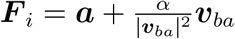. Now,

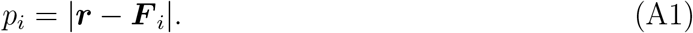

To obtain *q*_*i*_, note that *α* and *β* cannot be simultaneously negative. Then,

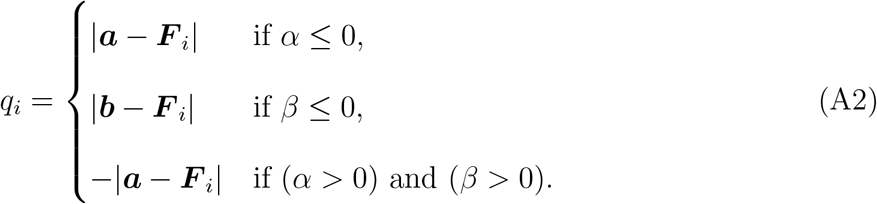

(The last line in the above equation can also be written as −|***b*** − ***F***_*i*_| owing to the oddness of the error function). With these and DT in place, it is then straightforward to obtain *K*(***r***) using Eq. (2).

## Appendix B: Codes

The scripts to generate the kernel density using EFNKDE are available for Python users at https://github.com/rahuljosephfernandez/EFNKDE_PY and R users at https://github.com/rahuljosephfernandez/EFNKDE_R.

Here, we have also included the Mathematica and Python codes, which define the function to calculate the Kernel density within the EFNKDE method. The last function (EFNKDE) in each of the code returns *K*(***r***) with bandwidth BW. “PointArr” is the array carrying the input data (spatial distribution of points). The same used in the codes below is only a place holder and does not correspond to the data used in this paper for generating any of the graphs. Also note that if the data is given in terms of latitudes and longitudes, then a conversion to 2D cartesian form is needed before using the below codes.

### 1. Mathematica Code for EFNKDE

~~~
PointArr = {{0.417, 0.429}, {0.422, 0.919}, {-0.694, 0.241}, {-0.718,0.515},
{-0.278, 0.156}, {-0.531, 0.004}, {0.337, 0.944}};
nPArr = Length[PointArr];
DelaunayTriangulatePlot = DelaunayMesh[PointArr];
AllEdgeList1 = MeshCells[DelaunayTriangulatePlot, 1][[All, 1]];
nAllEdgeList1 = Length[AllEdgeList1];
~~~

~~~
IntInfoGivenEdgePT[x_,y_,iEdge_] := Module[{PtX, ipta, iptb, Pta, Ptb, VXma, VXmb,
Vbma, VXmaDOTVbma, VXmbDOTVamb, FootOfThePerpendicular, LengthOfPerp, yA, yB},
PtX = {x, y};
ipta = AllEdgeList1[[iEdge, 1]];
iptb = AllEdgeList1[[iEdge, 2]];
Pta = PointArr[[ipta]];
Ptb = PointArr[[iptb]];
VXma = PtX - Pta;
VXmb = PtX - Ptb;
Vbma = Ptb - Pta;
VXmaDOTVbma = VXma. Vbma;
VXmbDOTVamb = VXmb. (-Vbma);
FootOfThePerpendicular = Pta + VXmaDOTVbma/Vbma. Vbma Vbma;
LengthOfPerp = Norm[PtX - FootOfThePerpendicular];
If[VXmaDOTVbma <= 0, yA = Norm[Pta - FootOfThePerpendicular]];
If[VXmbDOTVamb <= 0, yA = Norm[Ptb - FootOfThePerpendicular]];
If[(VXmaDOTVbma > 0) && (VXmbDOTVamb > 0),
yA = -Norm[Pta - FootOfThePerpendicular]];
yB = yA + Norm[Vbma];
{LengthOfPerp, yA, yB}
];
~~~

~~~
PartialEFNKDE[x_,y_,BW_,iEdgeEF_] := Module[{IntInfoval, sval, yAval, yBval,
LEdgeval, Expval, Denomval, erfsval, AnsvalEF},
IntInfoval = IntInfoGivenEdgePT[x, y, iEdgeEF];
sval = IntInfoval[[1]];
yAval = IntInfoval[[2]];
yBval = IntInfoval[[3]];
LEdgeval = yBval - yAval;
Expval = Exp[-(1/2) (sval/BW)^2];
Denomval = 2 Sqrt[2 \[Pi]] nAllEdgeList1 BW LEdgeval; erfsval = Erf[yBval/(Sqrt[2] BW)] - Erf[yAval/(Sqrt[2] BW)];
AnsvalEF = (Expval*erfsval)/Denomval;
AnsvalEF
];
~~~

~~~
EFNKDE[x_,y_,BW_] := Sum[PartialEFNKDE[x, y, BW, j], {j, 1, nAllEdgeList1}];
~~~

### 2. Python Code for EFNKDE

~~~
import numpy as np
import matplotlib.pyplot as plt
from scipy.spatial import Delaunay
import math
~~~

~~~
PointArr = np.array([[0.417,0.429], [0.422,0.919], [-0.694,0.241], [-0.718,0.515],
[-0.278,0.156], [-0.531,0.004], [0.337,0.944]])
~~~

~~~
tri = Delaunay(PointArr)
plt.triplot(PointArr[:,0], PointArr[:,1], tri.simplices)
ListOfTriangles = tri.simplices
~~~

~~~
def SortVertices(a,b):
  return [a,b] if a < b else [b,a]
~~~

~~~
AllEdgeList=[]
for i in ListOfTriangles:
  AllEdgeList.append(SortVertices(i[0],i[1]))
  AllEdgeList.append(SortVertices(i[1],i[2]))
  AllEdgeList.append(SortVertices(i[0],i[2]))
~~~

~~~
AllEdgeList1=np.unique(AllEdgeList, axis=0)
nAllEdgeList1 = len(AllEdgeList1)
~~~

~~~
def NormOfVec(V):
  return math.sqrt(np.dot(V,V))
~~~

~~~
def IntInfoGivenEdgePT(x,y,iEdge):
  PtX=(x,y)
  ipta = AllEdgeList1[iEdge,0]
  iptb = AllEdgeList1[iEdge,1]
  Pta = PointArr[ipta]
  Ptb = PointArr[iptb]
  VXma = PtX – Pta
  VXmb = PtX – Ptb
  Vbma = Ptb - Pta
  VXmaDOTVbma = np.dot(VXma,Vbma)
  VXmbDOTVamb = np.dot(VXmb,-Vbma)
  FootOfThePerpendicular = Pta + (VXmaDOTVbma/(np.dot(Vbma,Vbma)))*Vbma
  LengthOfPerp = NormOfVec(PtX - FootOfThePerpendicular)
  if VXmaDOTVbma <= 0.0:
    yA = NormOfVec(Pta - FootOfThePerpendicular)
  elif VXmbDOTVamb <= 0.0:
    yA = NormOfVec(Ptb - FootOfThePerpendicular)
  else:
    yA = -NormOfVec(Pta-FootOfThePerpendicular)
  yB = yA + NormOfVec(Vbma)
  return [LengthOfPerp,yA,yB]
~~~

~~~
def PartialEFNKDE(x,y,BW,iEdgeEF):
  IntInfoval = IntInfoGivenEdgePT(x,y,iEdgeEF)
  sval = IntInfoval[0]
  yAval = IntInfoval[1]
  yBval = IntInfoval[2]
  LEdgeval = yBval - yAval
  Expval = math.exp(−0.5*((sval/BW)**2))
  Denomval = 2*math.sqrt(2*math.pi)*nAllEdgeList1*BW*LEdgeval
  erfsval = math.erf(yBval/(BW*math.sqrt(2)))-math.erf(yAval/(BW*math.sqrt(2)))
  AnsvalEF = Expval * erfsval / Denomval
  return AnsvalEF
~~~

~~~
def EFNKDE(x,y,BW):
  Ans=0
  for iE in range(nAllEdgeList1):
    Ans = Ans + PartialEFNKDE(x,y,BW,iE)
  return Ans
~~~

## Notes

### Competing Interest Statement

The authors have declared no competing interest.

